# Extent of Molecular Chaperone Association Might Determine Fates of Membraneless Organelles during Aging in *C. elegans*

**DOI:** 10.1101/2021.12.17.473198

**Authors:** Pritam Mukherjee, Prajnadipta Panda, Prasad Kasturi

## Abstract

Proteome imbalance can lead to protein misfolding and aggregation which is associated with pathologies. Protein aggregation can also be an active, organized process and can be exploited by cells as a survival strategy. In adverse conditions, it is beneficial to deposit the proteins in a condensate rather degrading and resynthesizing. Membraneless organelles (MLOs) are biological condensates formed through liquid–liquid phase separation (LLPS), involving cellular components such as nucleic acids and proteins. LLPS is a regulated process, which when perturbed, can undergo a transition from a physiological liquid condensate to pathological solid-like protein aggregates.

To understand how the MLO-associated proteins (MLO-APs) behave during aging, we performed a comparative meta-analysis with age related proteome of *C. elegans*. We found that the MLO-APs are highly abundant throughout the lifespan. Interestingly, they are aggregating more in long-lived mutant worms compared to the age matched wildtype worms. GO term analysis revealed that the cell cycle and embryonic development are among the top enriched processes in addition to RNP components in insoluble proteome. Considering antagonistic pleotropic nature of these developmental genes and post mitotic status of *C. elegans*, we assume that these proteins phase transit during post development. As the organism ages, these MLO-APs either mature to become more insoluble or dissolve in uncontrolled manner. However, in the long-lived *daf-2* mutant worms, the MLOs may attain protective states due to extended availability and association of molecular chaperones.

## INTRODUCTION

Aging is a major risk factor for many diseases, including neurodegenerative, diabetes, cancer, and cardiovascular diseases (Hou et al. 2019). A major pathological hallmark of most of these disorders is protein misfolding and subsequent aggregation (Ross and Poirier, 2004) due to compromised protein homeostasis (proteostasis) (Hartl 2016). Proteostasis refers to the maintenance of the proteome in the correct conformation, concentration, and location required for cellular functions (Balch et al. 2008; Labbadia and Morimoto, 2015; Verma et al. 2021). Stress response and longevity pathways also promote proteostasis. For example, heat shock proteins (molecular chaperones) are expressed under various stress conditions to protect the proteome. They are targets of the conserved stress response pathways such as HSR (heat shock response) and IIS (insulin/IGF1-like signalling pathway) that affect proteostasis and life span (Hsu et al. 2003; Klaips et al. 2018). Indeed, it has been reported that long lived *daf-2* (insulin like receptor) mutant *C. elegans* activate a protective aggregation response during aging to maintain extended proteostasis (Walther et al. 2015). In part, this is achieved by inducing small heat shock proteins (sHSPs) expression, which in turn reduce the burden of misfolding proteins by sequestering them to aggregates or condensates. Notably, it has been observed that highly abundant proteins contribute more to the aggregate load due to their exceeding levels of solubility (super saturation). These supersaturated proteins form a metastable sub-proteome and undergo aggregation or interact with other aggregates during stress or aging. Supersaturation of the proteins is one of the driving forces for the increased protein aggregation (Ciryam et al. 2013, 2015, 2019; Walther et al. 2015).

Protein aggregates are considered inherently irreversible aberrant clumps, and are often associated with pathologies (Hipp et al. 2014; Soto and Pritzkow, 2018). However, recent evidence demonstrates that protein aggregation can also be an active, organized process (Fassler et al. 2021; Miller at al., 2015; Boronat et al. 2021). Physiological and reversible protein aggregation which seem to be beneficial has been observed in diverse systems from yeast to mammalian cells (Wallace et al. 2015; Saad et al. 2017; Becker and Gitler 2018; Cereghetti et al, 2021). Regulated and often reversible formation of protein aggregates or condensates perform a wide variety of cellular functions including response to stress, gene expression, cellular development and differentiation etc. (Alberti and Hyman., 2021; Fassler et al. 2021).

Recently, it has been demonstrated that under diverse conditions biological molecules can demix or remix (phase separation) with surrounding media. This phase separation in cells creates biomolecular condensate or membraneless organelles (MLOs) that can provide organization centre for biochemical reactions (Alberti, 2017; Boeynaems et al. 2018). Liquid-liquid phase-separation (LLPS) is a reversible molecular process that promotes the formation of biomolecular condensates or MLOs in living cells by proteins and RNA molecules, which have no fixed stoichiometry (Shin and Brangwynne 2017). For example, stress granules (SGs) are highly dynamic condensates formed in response to stress. These condensates are composed of two types of molecules, scaffolds and clients or regulators. While scaffold molecules have high number of valences and necessary for condensate formation, the client molecules have lower interaction valences and are recruited to condensates. These phase-separated and reversible assemblies provide a general regulatory mechanism as part of the stress response (Riback et al. 2017; Franzmann and Alberti 2019). In some situations, liquid to gel-like transition is essential cellular functions (Putnam et al. 2019; Bose et al. 2022). However, when this process is dysregulated, some condensates mature into non-dynamic, gel to solid like condensates, which are more toxic (Alberti and Carra 2018). These aberrant phase-transitions, might result in irreversible damage that acts as drivers of cellular aging (Alberti and Hyman 2016). It has been reported that the age dependent aggregated proteins have a significantly higher propensity for LLPS. Likewise, many supersaturated proteins also form condensates by phase-separation (Vecchi et al. 2020).

Organisms might utilize this fundamental mechanism of LLPS to cost-efficiently organize and reorganize cellular resources according to functional needs, especially during stress and aging (Franzmann and Alberti 2019; Alberti and Hyman., 2021). However, it is not yet clear how the cells select which form of condensates is preferred and beneficial for long term storage of biomolecules, and what regulates and maintains that preferred state of condensates from aberrant transitions. In an attempt to understand this, we choose to address the question of how the proteins capable of undergoing LLPS behave during aging and if aggregation of these proteins plays any cyto-protective roles by sequestration-based strategies. To do this, we obtained *C. elegans* MLO-APs from MLOS-MetaDb database (Orti et al. 2021) and investigated their behaviour during aging by comparing age associated proteomes of wildtype and long lived *daf-2* mutant worms and gonad less mutants (Walther et al. 2015; David et al. 2010). Our analysis revealed that majority of the MLO-APs maintain their higher abundance levels during the lifespan and sequester to aggregates or condensates. As the organism ages, these MLOs either mature to become more insoluble/solid-like or dissolve in uncontrolled manner. However, in the long-lived *daf-2* mutant worms, the MLOs may attain different protective states due to extended availability and association with molecular chaperones.

## METHODS

### Data Collection

Membraneless organelles associated proteins (MLO-APs) were downloaded from the MLOS-MetaDb (Orti et al. 2021) for *C. elegans* model organism. While downloading the dataset we used the all the options from PhasepPro, PhasepDB-Low Throughput, PhasepDB-High Throughput, DrLLPS-Scaffolds, DrLLPS-High Throughput, DrLLPS-Regulators. This resulted in a dataset, consisting of 413 protein entries with their LLPS associated characteristics (disorder content, LCRs, NLCRs as well as Sources, References).

Aging proteome of *C. elegans* was downloaded from the published datasets (Walther et al. 2015; David et al. 2010). Excel files containing age related changes in total and aggregate proteome of wildtype and *daf-2* mutant strains from Walther et al. (2015) were extracted and used for main analysis. To check for the gonad and germline less effects on the aggregation of MLO-APs, data from David et al. (2010) was used.

### Data Analysis

Proteome data was analysed using Perseus 1.6.15.0 (Tyanova et al. 2016), Microsoft Excel and GraphPad Prism 9 software. We first annotated the identified proteins based on categorical annotation provided by MLOS-MetaDb. We then cleaned our dataset based on this categorical annotation. Metascape server was used for analysing gene ontology and PPI networks. Graph Pad Prism 9 and ggplot2 in R (Wickham 2016) was used for graphical representation of data.

### Datasets

The datasets used in this work are available in the supplementary file.

## RESULTS

### Functional annotations of membraneless organelle associated proteins

To better understand the proteins that form biological condensates or MLOs, their function and regulation, we sought to identify their functional categories. For this, we downloaded known and predicted *C. elegans* specific MLO-APs from MLOS-MetaDb, a meta server for membraneless organelles associated proteins (Orti et al. 2021). This dataset consists of consolidated set of 413 proteins entries from three databases: PhaSePro (Mészáros et al. 2020), PhasepDB (You et al. 2020), DrLLPS (Ning et al. 2020) (supplementary table-1). We then looked for their biological processes by GO term analyses in Metascape (Zhou et al. 2019). All the *C. elegans* protein coding genes were considered for the background. In biological process category, we found that cell cycle (GO:0007049), translation (GO:0006412), embryo development (GO: 0009790), multi-organism reproductive process (GO:0044703), ribonucleoprotein complex biogenesis (GO: 0022613) were among the top enriched terms (Figure 1A).

**Figure 1.**
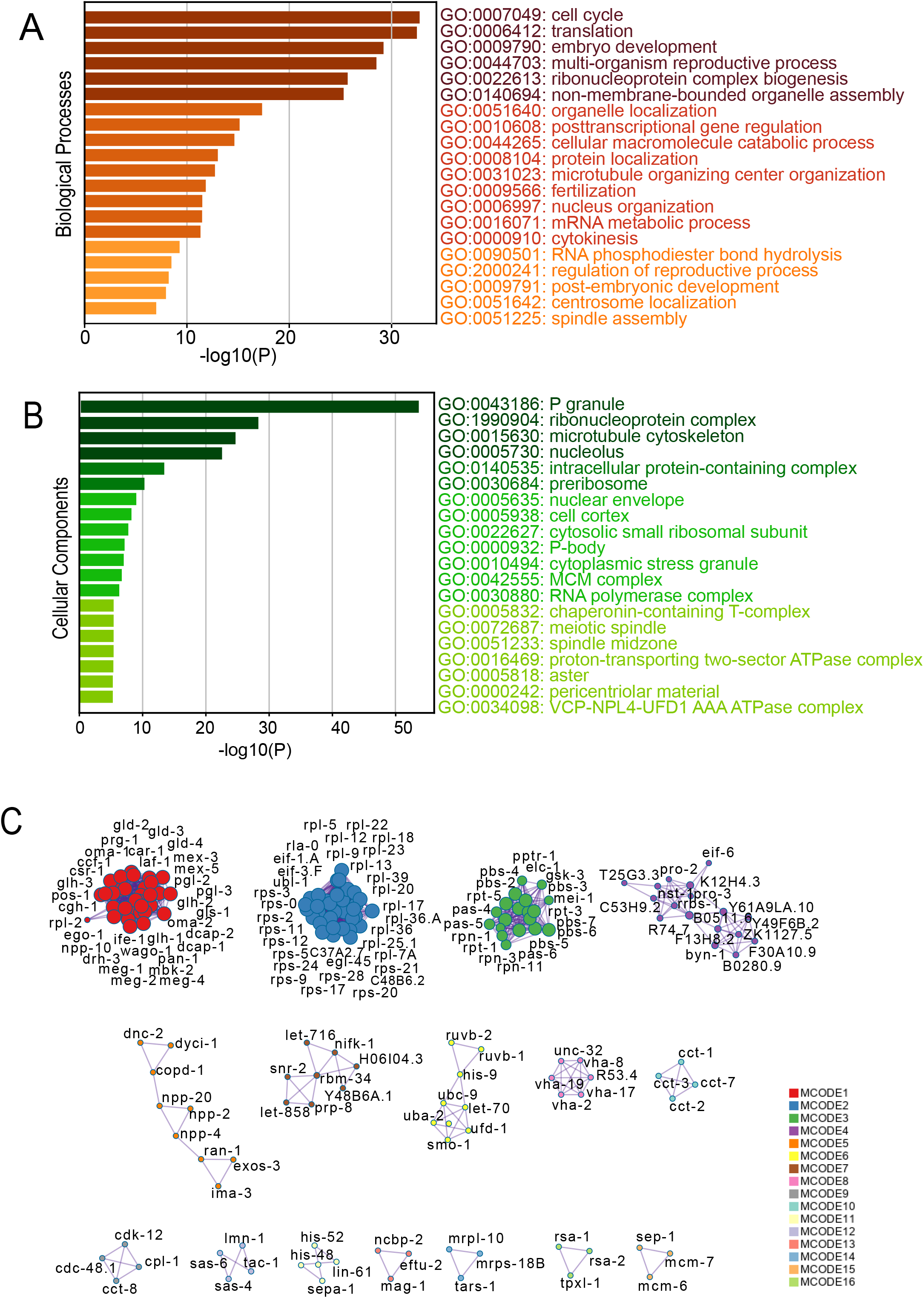
Functional annotation of MLO associated proteins. (A-B) Enriched gene ontology [GO] clusters of biological processes [BP] and cellular components [CC] of MLO associated proteins respectively (413 entries, Dataset from MLOS-MetaDb). (C) Protein-Protein Interaction networks of MLO associated proteins.

By doing GO cellular component analysis, we found the top enrichment in P granule (GO:0043186) followed by ribonucleoprotein (RNP) complex (GO: 1990904), microtubule cytoskeleton (GO:0015630), nucleolus (GO:0005730) (Figure 1B). We also checked the protein-protein interaction (PPI) network enrichment in Metascape, and found 16 Molecular Complex Detection (MCODE) networks (Figure 1C). The algorithm also provided the meanings by GO terms. We found the PPI enrichments for P-granule, m-RNP components, ribosomal subunit associated, proteasome, nucleolus, nuclear pore complex (NPC) and Chaperonin containing tailless complex polypeptide 1 (CCT) or tailless complex polypeptide 1 ring complex (TRiC). Interestingly we also found PPI enrichment of processes like cell cycle and developmental events. The network components and meanings are summarized in supplementary table-2. These results confirm that while some predicted MLO-APs are components of known biological condensates, others are yet to be identified as components of biological condensates or their role in this process.

### Total abundance levels of MLO associated proteins during aging

Age dependent decline in proteostasis contribute to the accumulation of misfolding proteins and also formation of aberrant, disease-causing condensates. In fact, there is a link between condensate-forming proteins and ageing-related diseases such as neurodegeneration (Alberti and Dormann 2019). When protein abundances exceed their solubility levels, they become supersaturated and form a metastable sub-proteome. Supersaturation of the proteins is one of the driving forces for this increased protein aggregation in human tissues and *C. elegans* (Ciryam et al. 2013, 2015, 2019; Walther et al. 2015).

To explore behaviour of the MLO-APs during aging, we investigated their abundance levels and aggregation by using published age-related proteomic dataset from wildtype and *daf-2* mutant worms (Walther et al. 2015). First, we looked for the abundance profiles of the MLO-APs that we analysed above. In total, 364 MLO-APs are identified in the aging proteome (supplementary table-3), however only 289 proteins are quantified in all the time points in the total proteome of both wildtype and *daf-2* mutant worms. These proteins are among the highly abundant proteins from day1 of adulthood and maintain their higher abundance throughout the lifespan (Figure 2A and 2B). Although they are among the highly abundant proteins, they are differentially expressed in wildtype and *daf-2* mutants as is evident from the heat map (Figure 2B). Interestingly, we observed that the granule forming propensity [catGRANULE] scores (Bolognesi et al. 2016) of these proteins do not correlate with their abundance intensities, that is the MLO-APs with highest granule forming propensity scores does not necessarily have highest LFQ intensity during aging (data not shown). It is unclear if the higher abundances of these MLO associated proteins are enough for them to undergo LLPS and form condensates, as their experimentally determined critical concentrations are not known. This suggest that intracellular milieu is the driving force for LLPS and condensate formation.

**Figure 2.**
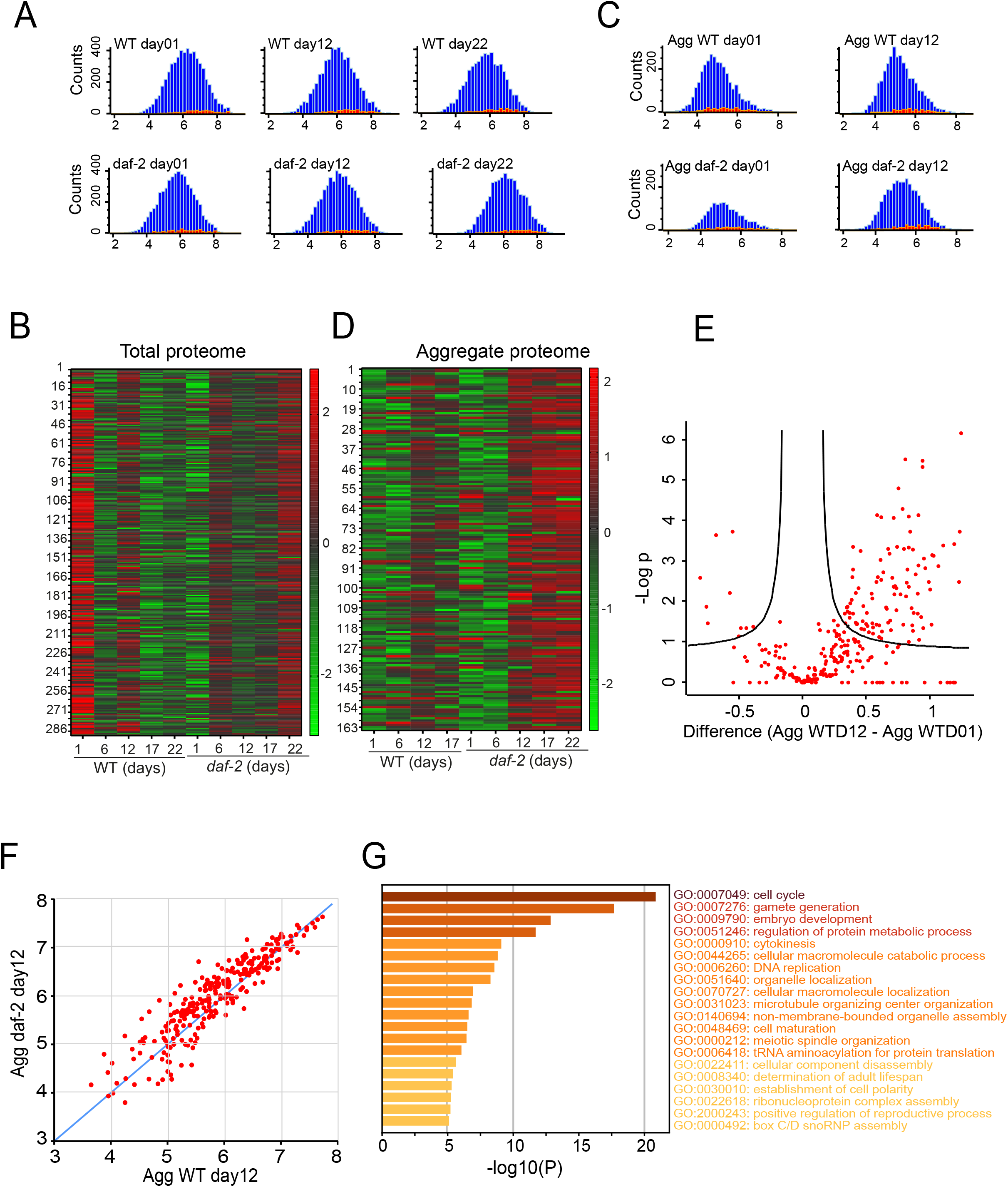
MLO associated proteins behaviour during aging. (A-B) Histograms showing the distribution of proteins in total proteome (A) and aggregate or insoluble proteome (B) identified during aging of *C. elegans*. Blue denotes the total number of proteins and red denotes the MLO associated proteins (364 entries). (C-D) Heat map showing the age and strain dependent z-score normalized LFQ intensity changes of MLO associated proteins in the total proteome (C) and aggregate or insoluble proteome (D). (E) Volcano plot showing the distribution of abundance levels of proteins between day1 and day12 of wildtype worms. (F) Scatter plot showing distribution of the same insoluble proteins in day12 wildtype and *daf-2* mutant worms (270 common entries, Pearson correlation coefficient 0.91). (G) Enriched gene ontology clusters of biological process of significantly upregulated MLO-APs in aggregate during aging.

### Aggregation of MLO associated proteins during aging

Membraneless organelles or biomolecular condensates are generally liquid like state, however with time they become more solid like. Although stress granules associated proteins become detergent insoluble with age (Lechler et al. 2017), it is unknown for other MLO-APs. Here we report that the 164 MLO-APs (quantified in all time points) that are identified in aggregated proteome in wildtype and *daf-2* mutant worms are weak detergent insoluble. We found that the aggregation of these proteins increases after day6 of adulthood as reported earlier (Walther et al. 2015) (Figure 2C and 2D). Our analysis shows that in wildtype worms out of 364 MLO-APs, 112 proteins are significantly upregulated and 10 proteins are downregulated in day 12 aggregate proteome (Figure 2E, supplementary table-4). Interestingly, the *daf-2* mutant worms contain more aggregation load compared to wildtype worms when considering common MLO-APs at day12 insoluble aggregated proteome (Figure 2F). Notably the upregulated insoluble proteins are enriched for the cell cycle (GO:0007049), gamete generation (GO:0007276), embryo development (GO: 0009790) regulation of protein metabolic process (GO: 0051246), cytokinesis (GO:0000910) and DNA replication (GO:0006260) (Figure 2G). Given the protective aggregation response of *daf-2* mutant worms and altered metabolism, we assume active reorganization of biological condensates including sequestration of other toxic / harmful molecules.

We then turned our attention to the analysis of proteins that are likely to become susceptible to aggregation when the control of protein homeostasis declines as in aging. Further we wanted to check if there is a commonality in wildtype and *daf-2* mutant worms aging associated sequestering of proteins in aggregates that functions in similar biological process. To answer this, we considered common proteins that have been quantified (at all the time points) in total and aggregate proteome of wildtype and *daf-2* mutant worms (wildtype total - 307, *daf-2* total - 298, wildtype aggregates - 210, *daf-2* aggregates - 170 proteins respectively (supplementary tables-5 and 6). We then performed GO term analysis and found that the enrichment of biological processes and cellular components are similar in both wildtype and *daf-2* mutant worms (Figure 3A and 3B). The following are among the top enriched terms for the proteins identified in aggregates: Translation (GO:0006412), cell cycle (GO:0007049), embryo development (GO:0009790) and embryo development ending in birth or egg hatching (GO:0009792) and DNA unwinding involved in DNA replication (GO:0006268). These results suggest possible roles for cell cycle and developmental proteins as MLOs.

**Figure 3.**
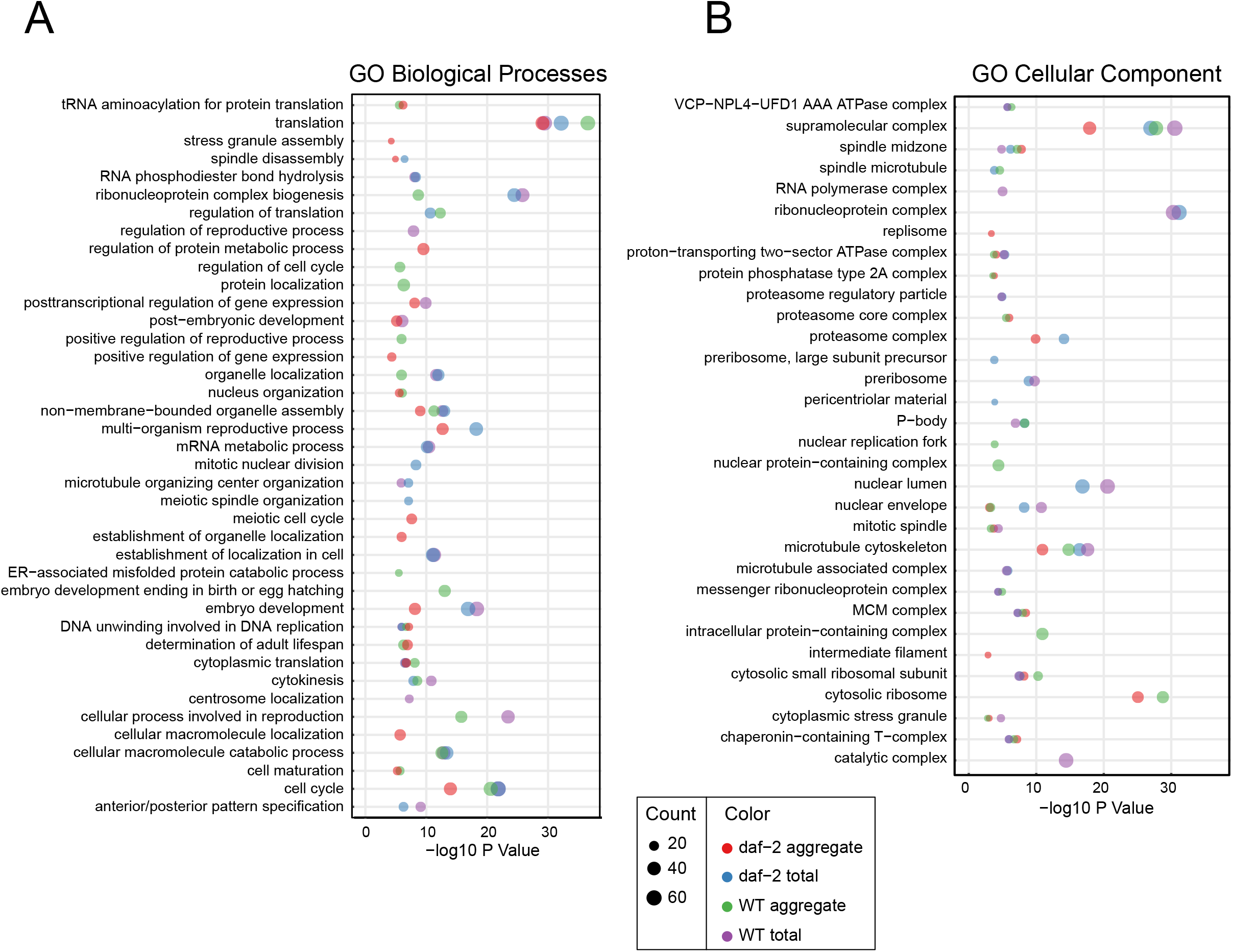
Age and strain dependent functional annotation of MLO associated proteins. (A) Enriched biological process of the common MLO associated proteins that have been quantified in all time points in total and aggregate proteome of wildtype and *daf-2* mutant worms during aging. (B) Enriched cellular components of the common MLO associated proteins that have been quantified in all time points in total and aggregate proteome of wildtype and *daf-2* mutant worms during aging.

### Age dependent behaviour of MLO associated cell cycle and developmental proteins

We found that the proteins associated with embryonic development, cell cycle and DNA replication are among the significantly enriched in aggregates. To get more insight on cell cycle (mitotic and meiotic) and developmental proteins during aging, we systematically checked fates of these proteins during aging in total and aggregated proteomes of wildtype and *daf-2* mutant worms (supplementary table-7).

When compared to day1 total proteome, there is no change in the abundance levels in wildtype worms, but in *daf-2* mutants there is a slight increase (Figure 4A, C and E). When compared to day1 aggregated proteome, there is an increase in the aggregate abundance levels both in wildtype and *daf-2* mutant worms. However, these protein aggregates are more enriched in *daf-2* mutant worms (Figure 4B, D and F).

**Figure 4.**
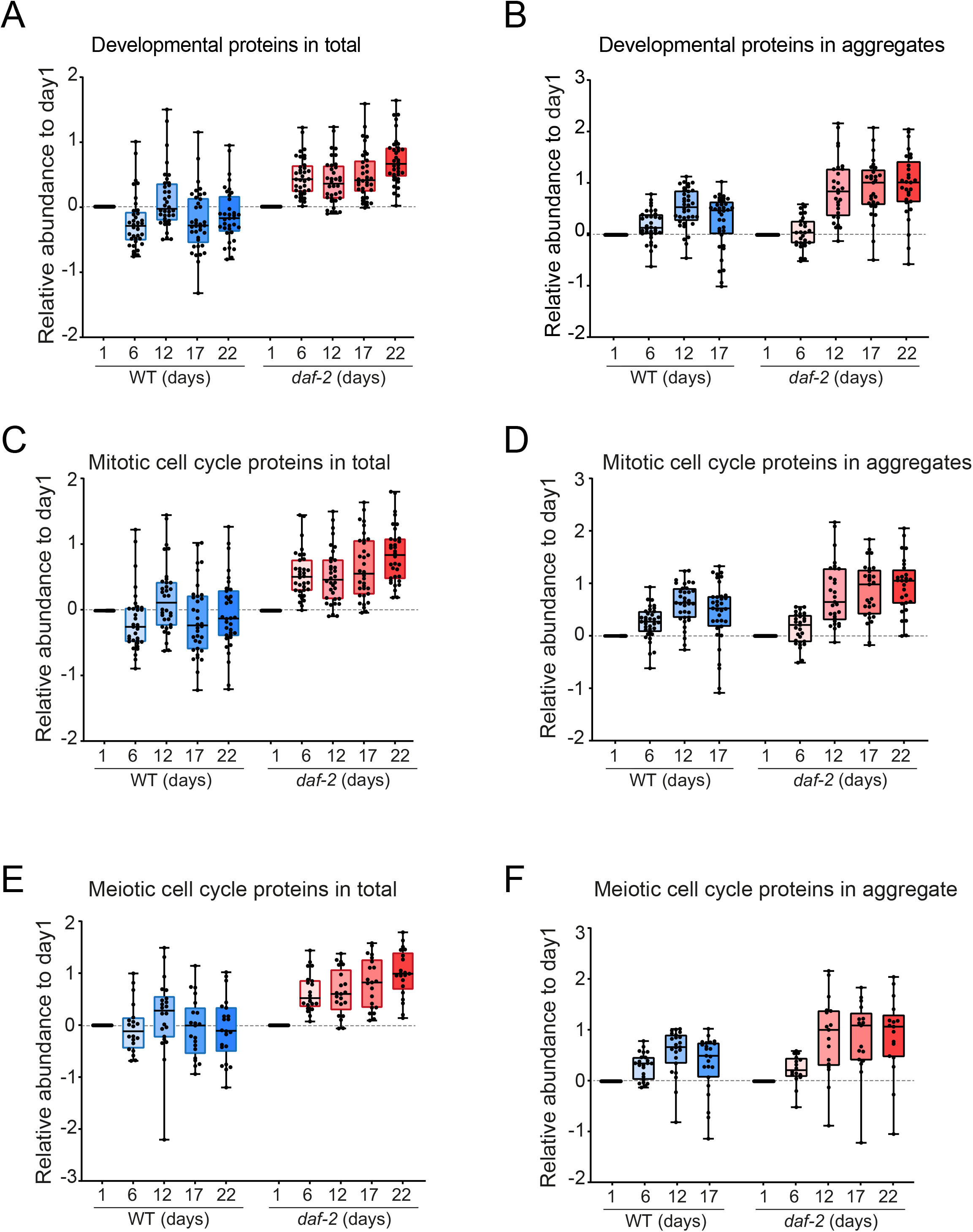
Relative abundance of MLO associated proteins involved in embryonic development and cell cycle (mitotic and meiotic) Box plots showing age and strain specific abundance changes (relative to day1) of proteins identified in total and aggregate proteomes. (A-B) Proteins involved in embryonic development (37 entries). (CD) Proteins involved in mitotic cell division (35 entries). (E-F) Proteins involved in meitotic cell division (24 entries).

Cell cycle and embryonic development related protein are enriched in the germline (Pu et al. 2017; Turner et al. 2015), suggesting a possibility that this protein aggregation is limited only to the germline. To rule out this possibility we checked other age dependent proteome aggregation data obtained from worm strains lacking gonad or germline (David et al. 2010). We found that embryonic developmental proteins and DNA replication related proteins are also shown to be aggregating during aging (supplementary table-8). Since, these are predicted as MLO-APs which are capable to undergo LLPS, we assume these proteins might be part of biological condensates or help in phase-transitions during aging.

### Age dependent behaviour of MLO associated PQC related components

Proper maintenance of proteostasis or the PQC machinery is essential for organismal survival and fitness especially during aging. Recent evidences suggest the extensive incorporation of proteostasis components into the dynamic assembly MLOs to spatiotemporally maintain their quality. Molecular chaperones are implicated in preventing the accumulation of misfolding proteins in condensates as well as aberrant phase transitions (Mateju et a., 2017; Gu et al. 2020, 2021; Boczek et al. 2021; Lu et al. 2021; Yoo et al. 2022), suggesting roles for PQC machinery in condensate quality maintenance.

Since aberrant condensate formation or disassembly are linked to protein aggregation diseases, we wanted to analyse PQC related components in the MLO-APs and how do they behave differently during aging in wildtype and long-lived *daf-2* mutant worms. From the MLOS-MetaDb, we found around 70 proteins as MLO-APs whose roles are ascertained to protein quality control including protein biogenesis, folding and degradation (supplementary table-7). Our analysis revealed that in wildtype worms, these proteins are relatively downregulated compared to day1 and are comparatively found more in aggregate fraction. In contrast, in *daf-2* mutant worms they are relatively upregulated compared to day1 and also enriched more in the aggregates compared to its age matched wildtype counterparts (Figure 5A and B). This again suggest existence of a protective response in *daf-2* mutans worms.

**Figure 5.**
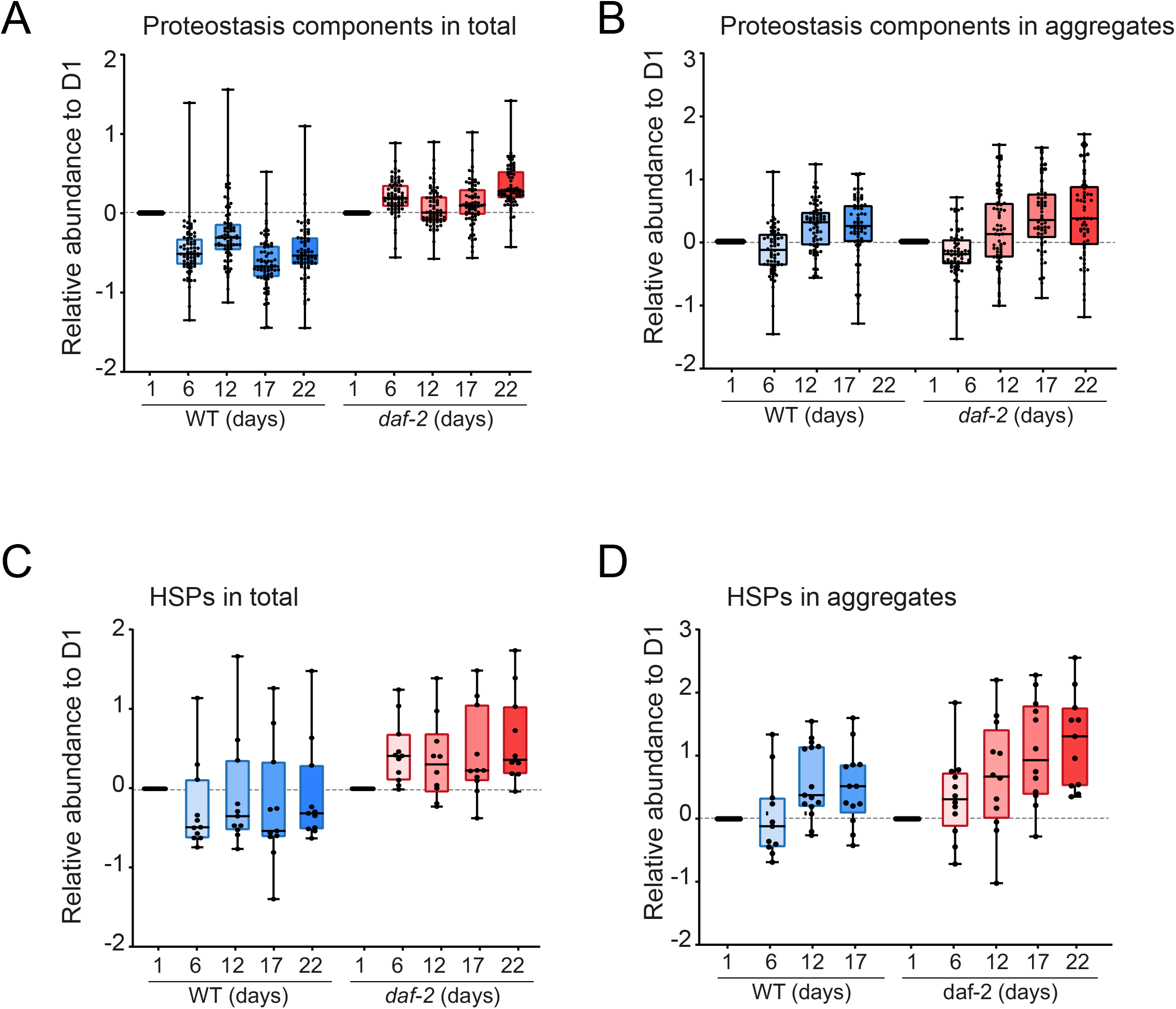
Relative abundance of MLO associated proteostasis components. Box plots showing age and strain specific abundance changes (relative to day1) of proteins identified in total and aggregate proteomes. (A-B) Proteostasis components excluding cytosolic HSPs HSP-70s and sHSPs (64 entries). (C-D) Cytosolic HSPs (HSP-70s and sHSPs 13 entries).

There have been reports about HSPs to be involved in the quality control of biomolecular condensates (Li and Leu 2022; Yoo et al. 2022). Therefore, we also looked separately on the cytosolic HSPs assuming they are also MLO-APs (which are not yet entered in MLOS-MetaDb dataset). We found that similar to other proteostasis component proteins, these HSPs are also upregulated in the total proteome of *daf-2* mutant worms compared to age matched wildtype worms. Likewise, these HSPs are also more strongly enriched in the aggregates of *daf-2* mutant worms. Among the HSPs, small heat shock proteins (sHSPs) are reported to promote sequestration of protein aggregation. This suggest that sHSPs may also actively promote formation of biological condensates. The altered metabolism in *daf-2* mutant worms may further act as buffer in modulating sHSPs as promoters of aggregation or phase-transitions of condensates. Indeed, it has been shown that overexpression of sHSPs results in beneficial effects. To further support, it has been shown that depletion of DAF-2 protein only late in life also result in lifespan extension (Venz et al. 2021). This strongly support, roles for PQC components modulating biological condensates for beneficial effects especially during aging.

## DISCUSSION

Membraneless organelles or biomolecular condensates are intracellular assemblies without surrounding a membrane that often form via liquid-liquid phase separation (LLPS). They have the ability to concentrate biomolecules and polymers to provide a spatial organization for biochemical reactions (Alberti 2017; Banani et al. 2017). Since condensates play key roles in cellular organization and physiology, their assembly and dissolution must be tightly controlled. Failure to control this regulation can result in defects from development to aging and can lead to diseases associated with protein aggregation. To better understand how MLO-APs behave during aging, we performed a comparative meta-analysis of predicted MLO-APs with aging proteome of *C.elegans*. Our analysis confirms that the predicted MLO-APs are components of known biomolecular condensates such as stress granules, p-bodies and nucleolus. In addition, identification of cell cycle and developmental proteins highlights importance of MLOs in cytoskeletal organization and cell division (Wiegand et al. 2020; Ong and Torres 2020). From the analysis, we assume that while some of these MLO-APs could act as scaffolds, majority of them are client or regulators.

Protein misfolding and aggregation are hallmarks of age-related diseases including neurodegeneration (Soto and Pritzkow, 2018; Hou et al. 2019). Aggregation-prone proteins are observed to be accumulated in the condensates and promote their hardening, indicating that both protein aggregation and condensation are probably linked. In fact, there is a link between condensate-forming proteins and ageing-related diseases such as neurodegeneration (Alberti and Dormann 2019). Interestingly, we found DNA replication, cell cycle and embryonic developmental related MLO-APs are significantly enriched in the aggregates. Abundance of these proteins increases during aging and exceed their solubility levels (supersaturated) resulting in formation of aggregates or aberrant phase transitions (if they form condensates during development). Why these proteins are more enriched in the insoluble fractions during aging? Why they are not removed post-development or in post-mitotic tissues?

Considering the nature of the analysed proteins as MLOs which can phase-transit, we assume two possible roles for these proteins getting sequestrated to condensates or aggregates during aging. First, during development these proteins might be sequestered to liquid like functional condensates and as the organism ages, these condensates gradually phase-transit to more solid like condensates. Finding these proteins in the aggregate fractions thus suggest a protective strategy for cells to sequester them from the pool so that their antagonistic pleotropic effects are reduced. Evolutionary theory of aging states that the developmental genes have antagonistic pleotropic nature. Their function is required during development; however, the same gene products might have negative consequences later in the age. Remarkably, it has been shown that reducing their levels by RNAi mediated knockdown during post developmental stages increases the life span of wildtype as well as *daf-2* mutant worms (Curran and Ruvkun 2007; Chen et al. 2007). Second, simply aggregating these proteins instead of degradation, further increase the burden on cellular proteostasis and may result in the possible detrimental effects. We assume the former case is true as the proteins required for developmental processes are also found to be aggregating during aging in worm strains lacking germline (David et al. 2010). It has been reported that organisms such as *C. elegans* and Xenopus begin life with an appreciable amount of amyloid material. However, their disassembly may lead to functional defects including developmental arrest. Indeed, it has been shown that ectopic expression of HSP104 (yeast disaggregase) in *C. elegans* caused cell division defects and embryonic lethality (Skuodas et al. 2020). These findings suggest that amyloid aggregates and/or biological condensates are a normal aspect of physiological functions and that their proper regulation is essential (Fassler et al. 2021). During aging the germline stem cells are reported to be quiescent, mitotic and meiotic processes comes to a halt and the proteins might not be properly distributed to the daughter cells. Thus, at that stage sequestering those proteins into aggregates or condensates but not degrading offer a protective strategy. If any divide signal is induced later in life (by sperm from a male), the proteins can readily be available for their function. It’s worth noting that disturbing p-granule condensates in the germline lead to trans-differentiation of germ cells to somatic cells (Updike et al. 2014), suggesting that the condensate organization is tightly regulated process.

Aberrant phase-transitions of biological condensates are linked to protein aggregation diseases. Recent evidences suggest roles for PQC machinery in condensate quality maintenance. Among the molecular chaperones, small heat shock proteins (sHSPs) are reported to promote sequestration of protein aggregation. Having multi-valent binding ability, sHSPs may act to seed and accumulate aggregate material in their near native confirmations for efficient refolding (Ungelenk et al. 2016; Mogk and Bukau 2017) and prevents aberrant phase-transitions (Boczek et al. 2021). This suggest that sHSPs might play roles in formation and maintenance of biological condensates.

There is compelling evidence that aging organism try to maintain their proteome balance by activating a proteome-scale protective aggregation response during aging. Sequestering of surplus or misfolded proteins into biomolecular condensates or MLOs is one such protective response during aging. This protective response is more pronounced in long lived *daf-2* mutant worms compared to wildtype (Walther et al. 2015). Membraneless organelles or biomolecular condensates are generally liquid like state, however with time they become more solid like. Given the protective aggregation response of *daf-2* mutant worms by increasing sHSPs levels and altered metabolism, we assume active reorganization of biological condensates during aging in these worms. This strongly support roles for molecular chaperones modulating biological condensates for beneficial effects especially during aging.

We conclude that during aging, the wildtype worms might sequester misfolding-prone proteins into condensates where the few MLO-APs act as scaffolds. However, with increasing age, these condensates due to insufficient PQC components, may aberrantly transit to toxic solids or abruptly dissolve their content. This leaking of the accumulated toxic molecular species might make inappropriate interactions with other molecules and eventually result in protostasis collapse. In contrast, *daf-2* mutant worms efficiently maintain condensate organization by recruiting more sHSPs and other factors to prevent or slow down aberrant phase-transitions and dissolution of the condensates. This extended availability and association with molecular chaperones in *daf-2* mutants eventually minimize leakage of misfolding proteins and prevent their inappropriate interactions with other molecules, which in turn reduce the burden on the PQC machinery. However, some of these assumptions and observations need to be experimentally verified.

## Supporting information

The datasets used in this work are available in the supplemental file

## Supplementary Information

The datasets used in this work are available in the supplemental file.

## Acknowledgements

This work was supported by SERB-CRG (Science and Engineering Research Board Core Research Grant, CRG/2021/007177 to P.K) and IIT Mandi seed grant (IITM/SG/PKS-78 to P.K). P.M acknowledges Ministry of Education and IIT Mandi for HTRA fellowship and P.P acknowledges DST for INSPIRE fellowship. The authors acknowledge BioX centre, SBS at IIT Mandi for providing facilities.

## Author contributions

P.M and P.K conceived the project. P.M performed analysis and made draft figures. P.P used R package to make figures. P.M, P.P and P.K analysed the data and performed the figure visualizations. P.M and P.K wrote the manuscript.

